# afpCOOL: An Accurate Tool for Antifreeze Protein Detection

**DOI:** 10.1101/231761

**Authors:** Morteza Eslami, Ramin Shirali-hossein-zade, Zeinab Takalloo, Ghasem Mahdevar, Abbasali Emamjomeh, Reza Hasan Sajedi, Javad Zahiri

**Affiliations:** Department of Computer Engineering, Arak University, Arak, Iran.; Computer Engineering Department, Sharif University of Technology, Tehran, Iran.; Department of Biochemistry, Faculty of Biological Sciences, Tarbiat Modares University, Tehran, Iran; Department of Mathematics, Faculty of Sciences, University of Isfahan, Isfahan, Iran; Laboratory of Computational Biotechnology and Bioinformatics (CBB), Department of Plant Breeding and Biotechnology (PBB), Faculty of Agriculture, University of Zabol, Zabol, Iran.; Bioinformatics and Computational Omics Lab (BioCOOL), Department of Biophysics, Faculty of Biological Sciences, Tarbiat Modares University, Tehran, Iran.; School of Biological Sciences, Institute for Research in Fundamental Sciences (IPM), P. O.Box 19395-5746, Tehran, Iran

**Keywords:** Antifreeze protein, Machine learning, Support vector machine (SVM), Physicochemical properties, Evolutionary profile

## Abstract

Various cold-adapted organisms produce antifreeze proteins (AFPs), which prevent to freeze of cell fluids by resisting the growth of the ice crystal. AFPs are currently being recognized in various organisms that are living in extremely low temperatures. AFPs have several important applications in increasing freeze tolerance of plants; maintain the tissue in frozen conditions and producing cold-hardy plants using transgenic technology. Substantial differences in the sequence and structure of the AFPs, pose a challenge for researcher to identify these proteins. In this paper, we proposed a novel method for identifying AFPs using support vector machine (SVM) by incorporating 4 types of features. Results on two benchmark datasets revealed the strength of the proposed method in AFP prediction. Also, according to the results on an independent test set, our method outperformed the current state-of-the-art methods. The further analysis showed the non-satisfactory performance of the BLAST in AFP detection: more than 62% of the BLAST searches have specificity less than 10% and there is no any BLAST search with sensitivity higher than 10%. These results reveal the urgent need for an accurate tool for AFP detection. In addition, the comparison results of the discrimination power of different feature types disclosed that evolutionary features and amino acid composition are the most contributing features in AFP detection. This method has been implemented as a stand-alone tool, namely afpCOOL, for various operating systems to predict AFPs with a user friendly graphical interface.

**Availability:** afpCOOL is freely available at http://bioinf.modares.ac.ir:8080/AFPCOOL/page/afpcool.isp

**Contact:** Dr Zahiri

## 1 INTRODUCTION

Organisms that are exposed to the freezing conditions produce special proteins, called antifreeze proteins (AFPs).(Commerou, et al., 2009; Graham, et al., 2013) AFPs bind to small ice crystals to forestall additional crystallization and depress the freezing point of solution below the melting point.(Davies, et al., 2002; Drori, et al., 2015; Drori, et al., 2015; Yeh and Feeney, 1996)The difference between the freezing and melting temperatures is referred to as thermal hysteresis. Therefore, additional terms have been proposed to naming the AFPs: ice structuring proteins and thermal hysteresis proteins.(Drori, et al., 2015; Venketesh and Dayananda, 2008)

For the first time, AFPs have been found in the species of fish and insect that have adapted to extremely low temperatures,(Davies, et al., 2002; Fletcher, et al., 1999) and the structure of 5 structurally distinct AFPs have been identified.(Davies and Hew, 1990) AFPs also have been found in fungi, bacterial species and overwintering plants.(Cheng, 1998; Ewart, et al., 1999; Logsdon and Doolittle, 1997) AFPs have potential applications in preservation, gene transformation, and cryosurgery of tumours and in agriculture for the production of economically resistance fishes and plants against extremely low temperatures.(ASSESSMENT, 2006; Fletcher, et al., 1999; Wang, et al., 1995)

Recently, two computational approaches have been proposed to predict AFPs,(Kandaswamy, et al., 2011; Zhao, et al., 2012) but these methods do not have a satisfactory performance. Considering the substantial differences in the sequence and structure of AFPs,(Griffith and Yaish, 2004; Jia and Davies, 2002) it is need to use suitable machine learning methods to predict AFPs. An appropriate machine learning algorithm offers a cost effective approach to construct predictive model to identify AFPs by exploiting experimentally validated training data. In this article, we proposed a computational method to identify AFPs. Our method achieved accuracy of 95% and 91% on two benchmark datasets and accuracy of 96% on an independent test dataset.

## 2 METHODS

### 2.1 Dataset

In this study, we used two independent datasets (Fig. 1) to compare prediction performance of the proposed method with the current state-of-the-art methods and to evaluate the strength of the predictor.

**Fig.1.**
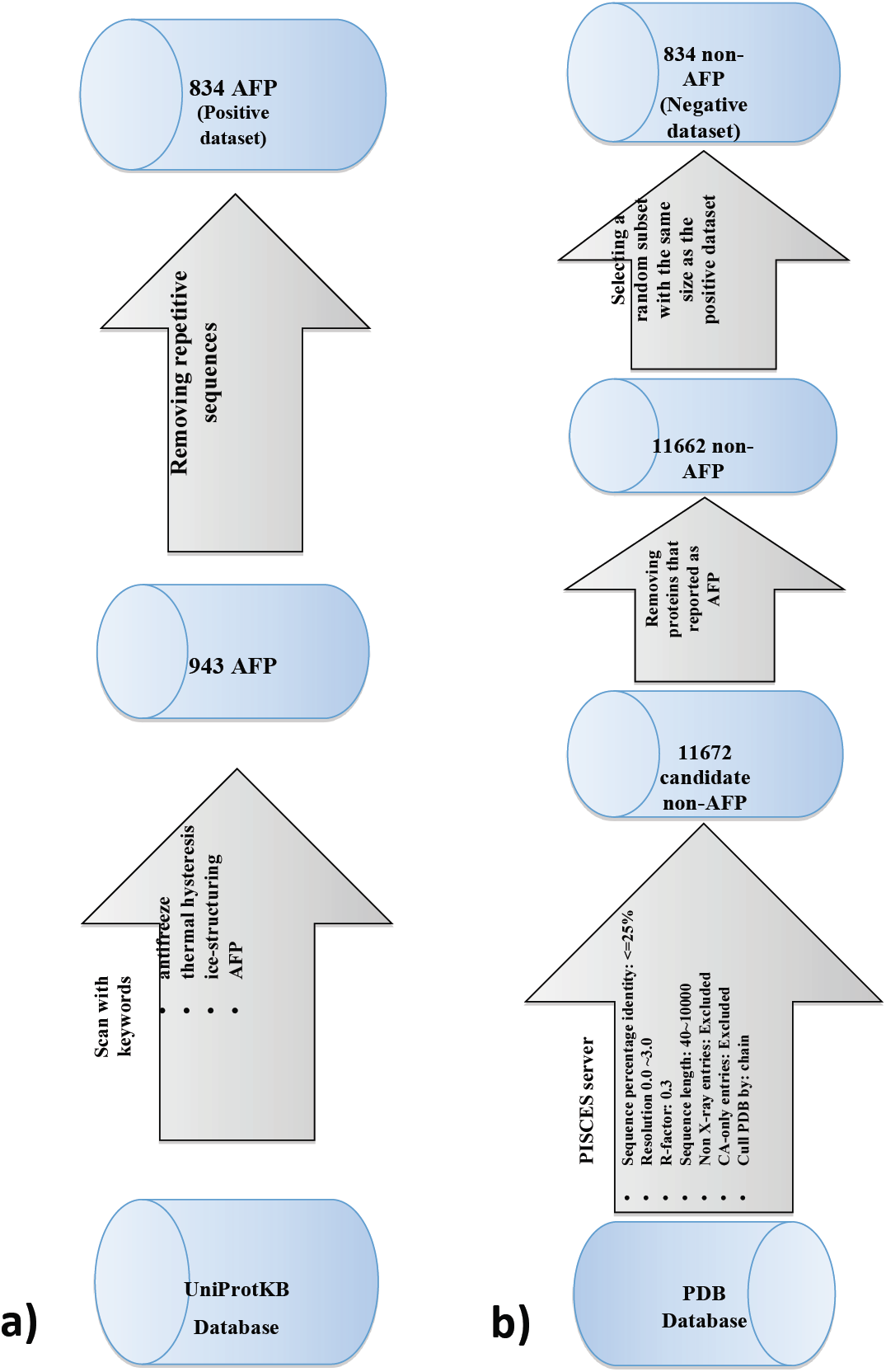
Schematic representation of AFP834 dataset construction. a) Positive dataset contains 843 AFPs. b) Negative dataset contains 843 non-AFPs.

#### 2.1.1 AFP481 Dataset

The first benchmark dataset (AFP481) has been retrieved from Kandaswamy et al.(Kandaswamy, et al., 2011) This dataset contains 481 AFPs and 9493 non-AFPs that have been used for the positive and the negative dataset, respectively. In this dataset, the proteins with >=40% sequence similarity were omitted using CD-HIT.(Li, et al., 2001) Train dataset consists of 300 AFPs out of all the 481 AFPs and 300 non-AFPs out of all the 9493 non-AFPs that have been selected randomly for positive and negative datasets, respectively. Also, remaining 181 AFPs and 9193 non-AFPs have been used as an independent test dataset.

#### 2.1.2 AFP834 Dataset

To have a better assessment of strength of the proposed method, a more comprehensive benchmark dataset, namely AFP834, has been assembled as follows (Fig. 1b). Antifreeze protein sequences were retrieved from the UniProtKB database.(Bairoch, et al., 2005) For this goal, UniProtKB has been scanned with a list of keywords that implying antifreeze proteins. The used keywords are: “antifreeze”, “thermal hysteresis”, “ice-structuring” and “AFP”. We retrieved 943 proteins in total, which the number of proteins corresponding to “antifreeze”, “thermal hysteresis”, “ice-structuring” and “AFP” keywords are: 734, 22, 52 and 135, respectively. Finally, after removing 109 repetitive proteins, the dataset had contained 834 non-redundant AFPs.

To select negative examples (non-AFP proteins), we took advantage of PISCES(Wang and Dunbrack, 2003) that is a public server for culling sets of protein sequences from the Protein Data Bank (PDB)(Berman, et al., 2000)by sequence identity and structural quality criteria (Fig. 1). We used the same number of non-AFP proteins as the number of AFP proteins to construct a balanced dataset.

### 2.2 Features

We trained our model to detect AFPs by exploiting four types of descriptors: hydropathy (3 descriptors), physicochemical properties (218 descriptors), amino acid composition (20 descriptors) and evolutionary profile (400 descriptors).

#### 2.2.1 Hydropathy descriptors

According to the hydropathy, 20 amino acids were categorized to 3 feature groups as follow:strongly hydrophilic (RDENQKH); strongly hydrophobic (LIVAMF) and weakly hydrophilic or weakly hydrophobic (STYW). For every feature of these three groups in a protein sequence, the number of occurrences of each group has been computed and divided by the length of the sequence.

#### 2.2.2 Physicochemical descriptors

To compute physicochemical descriptors, 544 different physicochemical indices have been extracted from AAINDEX(Kawashima and Kanehisa, 2000) database, which is a database of numerical indices representing various physicochemical and biochemical properties of amino acids. To reduce the biases, which can be result of having many correlated indices, we considered any pair of indices with correlation coefficient greater than 0.8 and less than -0.8 as redundant indices. And finally, a subset of 218 non-redundant indices has been selected to encode proteins.

#### 2.2.3 Amino acid composition

Each protein has been encoded by a 20-dimentional feature vector that shows the amino acid composition of the protein. Every element of this vector is the frequency of an amino acid in the protein sequence.

#### 2.2.4 Evolutionary information

Evolutionary information has been shown to be effective to detect diverse properties of proteins.(Zahiri, et al., 2014; Zahiri, et al., 2013; Zhao, et al., 2012) We used evolutionary information of a protein in the form of Position-Specific Scoring Matrix (PSSM). To generate PSSMs for all the proteins, Position Specific Iterated BLAST (PSI-BLAST) has been used against the NCBI non-redundant dataset with three iterations and the e-value of 0.0001. Regarding substitution scores in the PSSMs, each protein has been encoded as a 400-dimentional feature vector. Each element of this vector is the sum of all positive substitution scores of a specific amino acid to the one of the twenty standard amino acids.

#### 2.2.5 Normalization

To avoid the bias, which can be result of different sequence length, in all four above mentioned descriptor types every element has been normalized by the length of the sequence.

### 2.3 Support Vector Machines

In recent years, support vector machine (SVM) has been widely used in various prediction problems in bioinformatics.(Zahiri, et al., 2013) SVM classifies the input samples, represented in the form of n-dimensional feature vectors, into two classes using an optimal hyper-plane in the feature space. In this study, the input to the SVM classifier is a 644-dimensional vector that encodes the features of a given protein, and, the output is a binary label indicating whether the given protein is an AFP or not. We used the SVM implementation in the WEKA package(Hall, et al., 2009) with Pearson VII function-based universal kernel (PUK).

### 2.4 Evaluation parameters for the prediction performance

A 10 fold cross-validation approach was used to evaluate the performance of the proposed prediction model. Positive and negative instances were distributed randomly into 10 folds. In each of the 10 iterative steps, 9 of the 10 folds were used to train the classifier, and then the classifier was evaluated using the remaining data (test data). The predictions made for the test instances in all the 10 iterations were combined and used to compute the following performance measures:

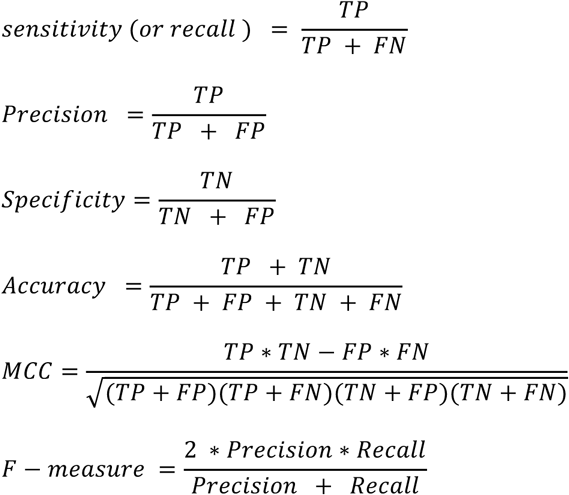

Where, TP and TN are the correctly predicted AFP and non-AFP instances, respectively. Similarly, FP and FN are the number of proteins that wrongly predicted as AFP and non-AFP, respectively. In addition to the above mentioned measures, we used the receiver operating characteristic (ROC) curve, which is an important graphical tool for assessing the classification performance. ROC plots sensitivity (true positive rate) against false positive rate and shows the trade-off between sensitivity and specificity. We also, used the area under the ROC curve (AUC), as a reliable performance measure.

## 3 RESULTS & DISCUSSION

To show the strength of the proposed regarding the current state-of-the-art methods, we compared afpCOOL with two recently published methods: AFP-Pred^17^and AFP-PSSM.(Zhao, et al., 2012) The prediction results of afpCOOL on the two benchmark datasets, AFP481and AFP834, and an independent test set have been described in the following.

### 3.1 Results on the AFP481 dataset

Table 1 shows the most important prediction performance measures for afpCOOL on the AFP481 dataset. This dataset contains 300 AFPs proteins and 300 non-AFPs. According to the 10-fold cross validation results, our method performed well in AFP detection: the method achieved an accuracy of 93% with an f-measure of 93%.

**Table 1.** Prediction performance of the afpCOOL on two benchmark datasets in a 10-fold cross validation procedure. The AFP481 dataset contains 300 AFPs and 300 non-AFPs; and, the AFP834 dataset contains 834 AFPs and 834 non-AFPs.

| Performance measures \ Datasets | Accuracy | Precision | Sensitivity | F-Measure | MCC | AUC |
| --- | --- | --- | --- | --- | --- | --- |
| **AFP481** | 0.93 | 0.93 | 0.93 | 0.93 | 0.87 | 0.93 |
| **AFP834** | 0.91 | 0.92 | 0.92 | 0.92 | 0.84 | 0.92 |

### 3.2 Results on the AFP834 dataset

This dataset contains 1668 proteins with equal number of positive (AFPs) and negative (non-AFPs) instances. The various performance measures in a 10-fold cross validation procedure revealed the strength of afpCOOL (Table 1). Our method achieved an accuracy of 91.9%, precision and sensitivity of 92% with an MCC of 84%. Also, Fig. 2 shows the ROC curve of the method on the two mentioned benchmark datasets. As these curves disclose, the proposed model performed better on the larger dataset (AFP834 dataset).

**Fig. 2.**
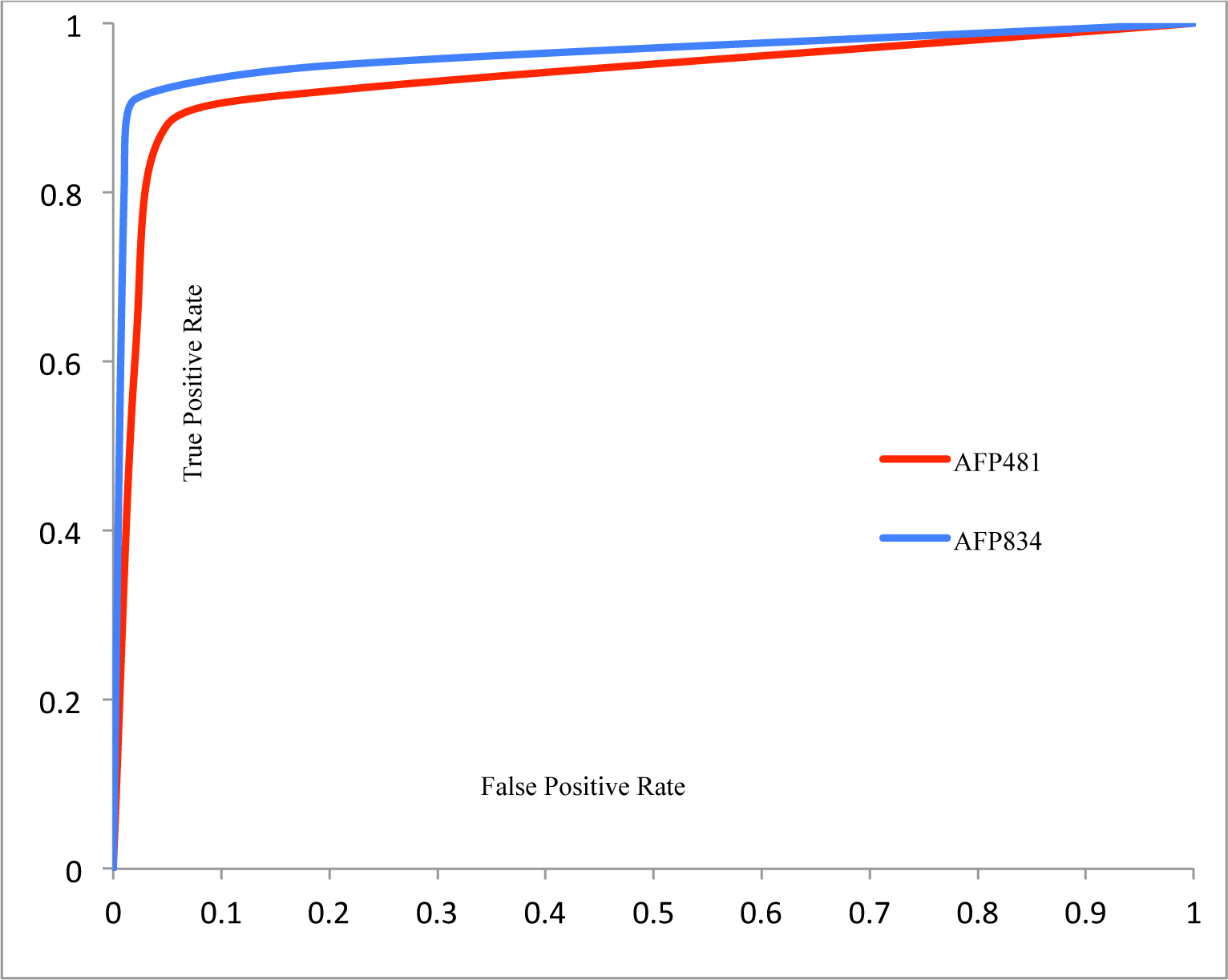
The receiver operating characteristic (ROC) curves of our method on the two benchmark dataset calculated from the ten-fold cross validation. The AFP481 dataset contains 300 AFPs and 300 non-AFPs; and, the AFP834 dataset contains 834 AFPs and 834 non-AFPs.

### 3.3 Comparison with the current state-of-the-art methods

For further evaluation of our method, an independent test set has been used to compare the method with the current state-of-the-art methods. We used a test dataset that recently exploited by Zhao et al(Zhao, et al., 2012) as the benchmark for comparison purpose, which contains 181 AFPs and 9193 non-AFPs. To have a fair comparison, we have trained our model with the same data that has been used by the two competitor methods (AFP481).

As Table 2 shows, AFP-Pred performed better in sensitivity measure, but is the worse method according to the specificity and accuracy.

**Table 2.** Performance comparison of the proposed method (afpCOOL) with the two current state-of-the-art methods in AFP prediction. All methods are trained on the AFP481 dataset in a 10-fold cross validation procedure and tested on an independent test dataset with 181 AFP and 9193 non-AFPs.

| Performance Measure<br>Method |  |  |  |
| --- | --- | --- | --- |
|  | Sensitivity | Specificity | Accuracy |
| **afpCOOL** | 0.72 | 0.98 | 0.96 |
| **AFP-Pred**(Griffith and Yaish 2004) | 0.85 | 0.82 | 0.83 |
| **AFP-PSSM**(Zhao et al. 2012) | 0.76 | 0.93 | 0.93 |

The accuracy of afpCOOL (96%) is higher than the accuracy of AFP-Pred (83%) and accuracy of AFP-PSSM (93%). Also, our method outperformed the other methods in terms of specificity.

### 3.4 afpCOOL tool

We have developed the afpCOOL as a tool that enables fast *in-silico* AFP detection. This tool has been implemented as a stand-alone java application for various operating systems with a user friendly graphical interface (Fig. 3). Users can use the afpCOOL simply by providing sequences (in fasta format) and PSSMs of the interested proteins; afpCOOL extracts features from the provided inputs and then uses the trained SVM-based model to assign AFP or non-AFP label to the queries. The afpCOOL is freely downloadable for non-commercial use at http://biocool.ir.

**Fig. 3.**
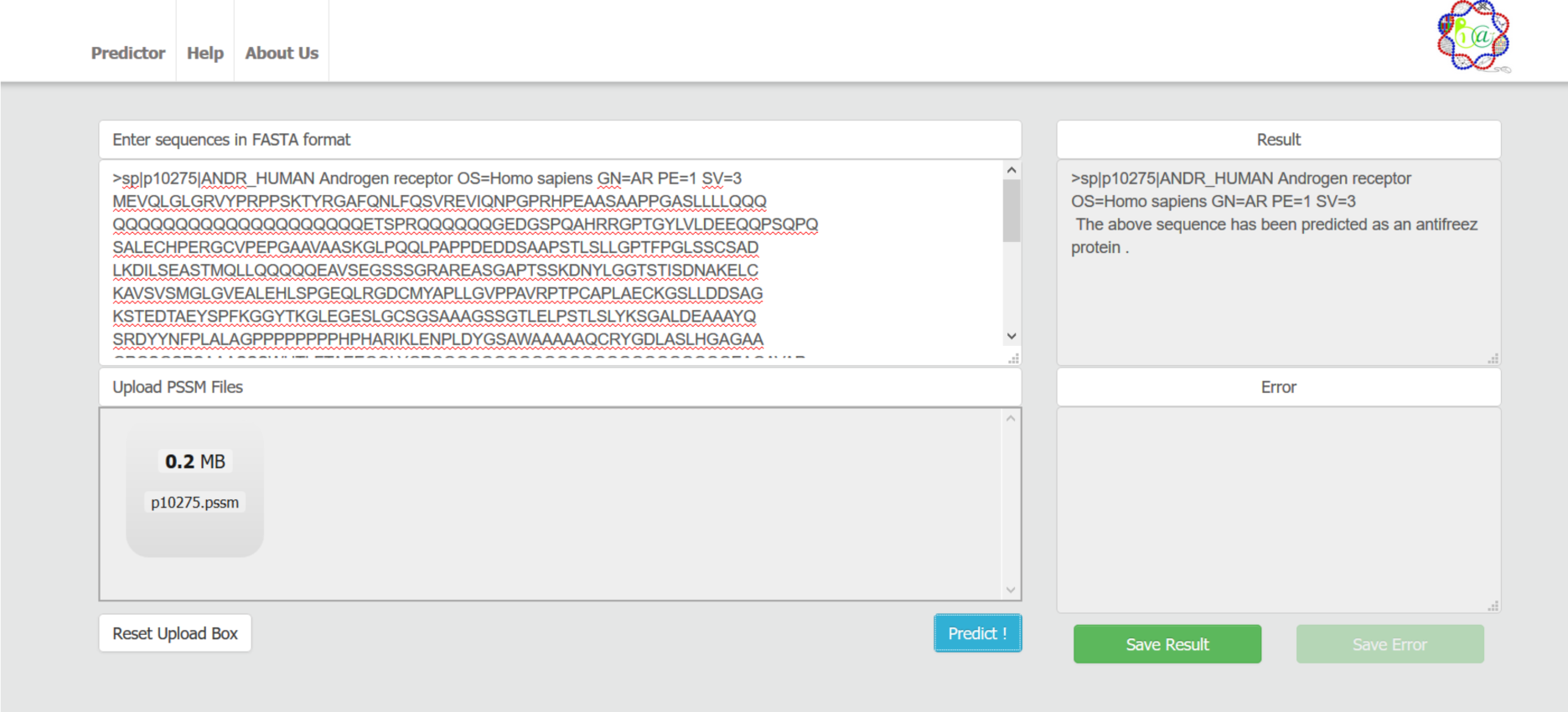
Graphical user interface of afpCOOL.

### 3.5 The urgent need for a reliable computational tool for AFP prediction: BLAST cannot effectively detect the AFPs

To show the strength of the afpCOOL, we have run BLAST when using each of the AFP as the query against the last update (August 12, 2015) of the UniProt Archive (UniParc) database with e-value<=1e-3. It should be mentioned that 801 proteins have been mapped to UniParc identifiers, and as the result, 801 BLAST searches have been used for this analysis. As Supplementary Fig. S1a shows, the BLAST does not have a satisfactory specificity in AFP detection: more than 62% of the BLAST searches (539 out of 801) have specificity less than 10%; also, more than 87% of the BLAST searches have specificity <=50%. Considering the large dataset that used for BLAST searches, it may be expected to have a low specificity and very good sensitivity. Nevertheless, considering the sensitivity of the BLAST searches the story is worse than the specificity (Supplementary Fig. S1b): there is no any BLAST search with sensitivity higher than 10%; also, 84% of the BLAST searches have sensitivity <=5% (673 search out of 801 BLAST searches).

Interestingly, there are 35 proteins with no any search results in the BLAST search (Supplementary Table S1) and 11 proteins with 0% sensitivity (Supplementary Table S1). As we can see, 5 out of 11 (45%) proteins with 0% sensitivity extracted from Boreogadus saida (Polar cod). In addition, the proteins of these two categories have a significantly different amino acid composition regarding the other AFPs (Supplementary Fig. S2). As it is shown in the Supplementary Fig. S2, alanine is the major building blocks of these AFPs (43% in the AFPs with no any hit in the BLAST search and 47% in the AFPs with 0% sensitivity in the BLAST search). It can be due to the abundant alpha helix in the structure of these proteins: there is only one structure for P04367 (PDB ID: 1Y03) that confirms this hypothesis.

### 3.6 Discrimination power of the different feature types: evolutionary features are the most contributing features in AFP detection

To assess the discrimination power of the four different feature types, we have re-trained the SVM model using each feature types when discarding the other. As Fig. 4 shows, evolutionary features (PSSM-based features) are the most discriminant feature in AFP detection: this feature type has the best performance in the all six performance measures. Also, as it may be expected, amino acid composition is the second most contributing feature type in AFP detection.

**Fig. 4.**
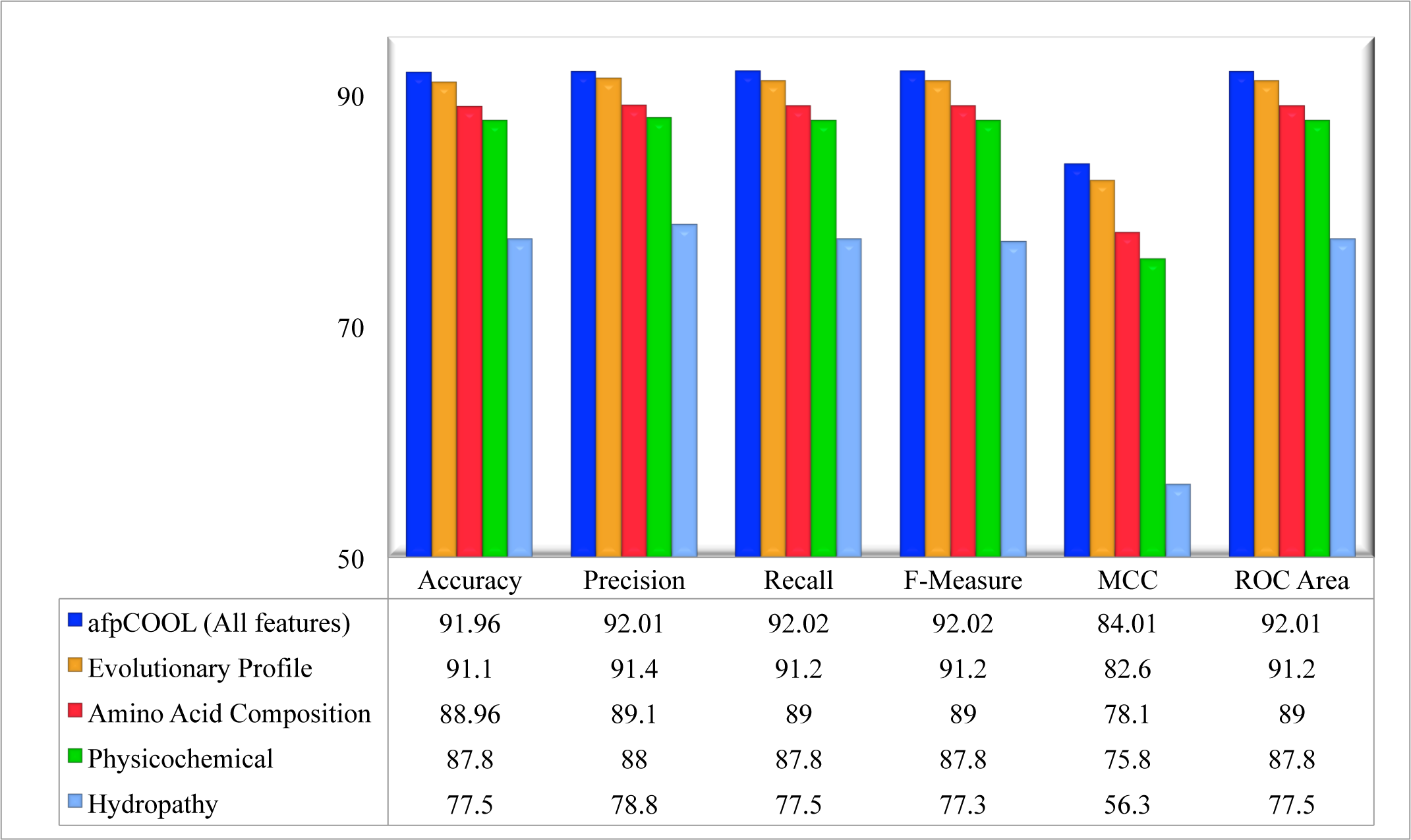
Discrimination power of the four different feature types, when the SVM model has been re-trained using each feature types while discarding the other.

## 4 CONCLUSION

We developed a novel SVM-based method to predict AFPs. In this method, each protein has been encoded by four features (evolutionary profile, amino acid composition, Hydropathy and physicochemical properties). The results showed that these types of features can significantly improve the prediction of AFPs. The obtained results on two benchmark datasets revealed the strength of our method in AFP detection. Also, the results on an independent test set confirmed the better performance of the proposed method regarding the current state-of-the-art AFP prediction methods. In addition, our more analysis disclosed the poor performance of BLAST in AFP detection and so indicates the critical need to an accurate tool for this purpose. Finally, evolutionary profile and amino acid composition showed the most power in discriminating AFPs from non-AFPs.

